# Metagenomes and Metagenome-Assembled Genomes from Tidal Lagoons at a New York City Waterfront Park

**DOI:** 10.1101/2025.05.01.651762

**Authors:** Sally Kong, Eliana Abrams, Yehuda Binik, Christina Cappelli, Mathew Chu, Taiyo Cornett, Isayah Culbertson, Epifania Garcia, Jada Henry, Kristy Lam, DB Lampman, Grace Morenko, Illusion Rivera, Tanasia Swift, Isabella Torres, Rayven Velez, Elliot Waxman, Serena Wessely, Anthony Yuen, Casey K. Lardner, JL Weissman

## Abstract

We report the sequencing and analysis of 15 shotgun metagenomes, including the reconstruction of 129 high-quality metagenome-assembled genomes, from tidal lagoons and bay water at Bush Terminal Piers Park in Brooklyn, NY sampled from July to September 2024.. Our metagenomic database for this site provides an important baseline for ongoing studies of the microbial communities of public parks and waterfront areas. In particular, we provide rich functional and taxonomic annotations that enable the use of these metagenomes and metagenome-assembled genomes for a wide variety of downstream applications.

## Introduction

We report the sequencing and analysis of 15 shotgun metagenomes from tidal lagoons and bay water at Bush Terminal Piers Park in Brooklyn, NY from July to September 2024. Notably, this waterfront park is an active site of ecological research and restoration by the Billion Oyster Project, an environmental nonprofit whose mission is to restore oyster reefs to New York Harbor through public education initiatives. Billion Oyster Project maintains an active community oyster reef in the innermost of our focal lagoons [2, 3].

Bush Terminal Piers Park is developed on a former brownfield, subject to storm- and sea level rise-related flooding, and is a social and environmental amenity for area residents. In combination with efforts to rezone nearby industrial areas for mixed-use development, the area is also impacted by the contested forces of gentrification [1]. In addition to sports fields, there are a series of short nature trails through a small, wooded area, and walking paths near our focal lagoons. These lagoons have been used for community events (e.g., a community boating event in summer 2024) and we frequently encountered members of the public using these spaces.

In aquatic ecosystems, bivalve populations exert strong top-down control on microbial communities via size-dependent predation of larger microbes [4] and simultaneously redirect nutrients back to these communities through their excretions which are in-turn remineralized by microbes [5, 6], making their impact on community structure hard to predict. Second order effects of bivalve addition, including changes to local hydrology and sedimentation rates, further complicate this picture [5]. These effects may in turn potentially feedback on oyster population health. In short, it is difficult to predict how the restoration of oyster reefs around New York Harbor will alter local microbial community structure and function. Complicating things further, we do not have a detailed baseline for the microbial community at reef-impacted sites. We constructed a metagenomic time series at this site during mid-to-late summer of 2024 in order to build a location-specific database that will serve as an important resource for future studies of the microbial populations in NYC’s waters, particularly at sites of active restoration like Bush Terminal Piers.

## Materials & Methods

### Sample Collection

Water samples were collected on four sampling dates from July to September 2024 at Bush Terminal Piers Park, Brooklyn, NY, during low tide, when the site forms distinct inner and outer lagoons disconnected from the bay, with the oyster reef located in the inner lagoon (Table 1). On the day of collection, water samples were immediately vacuum-filtered onto 0.22 µm Cellulose Nitrate Filter membranes (Sigma Aldrich GSWP04700). The filter membranes were then stored at -80°C until DNA extraction.

**Table 1.** Sample details

| Sample | Site | Date | Water Temperature (C) | Notes |
| --- | --- | --- | --- | --- |
| A01 | Inner Lagoon (IL) | 07-19-24 | 28.4 |  |
| A02 | Inner Lagoon (IL) | 07-19-24 |  |  |
| A03 | Inner Lagoon (IL) | 08-05-24 | 25.0 |  |
| A04 | Inner Lagoon (IL) | 08-05-24 |  |  |
| A05 | Inner Lagoon (IL) | 08-05-24 |  |  |
| A06 | Bay Water (R) | 09-02-24 | 24.0 |  |
| A07 | Inner Lagoon (IL) | 09-02-24 | 24.3 |  |
| A08 | Inner Lagoon (IL) | 09-02-24 |  |  |
| A09 | Outer Lagoon (OL) | 09-02-24 | 24.1 |  |
| A10 | Outer Lagoon (OL) | 09-02-24 |  |  |
| A11 | Inner Lagoon (IL) | 09-17-24 | 24.3 |  |
| A12 | Inner Lagoon (IL) | 09-17-24 |  |  |
| A13 | Outer Lagoon (OL) | 09-17-24 | 24.6 |  |
| A14 | Outer Lagoon (OL) | 09-17-24 |  |  |

### DNA Extraction and Sequencing

DNA was extracted from the stored filters using the Qiagen DNEasy PowerWater Kit (14900-100-N) following the manufacturer’s protocol. Extracted DNA was quantified using the Qubit dsDNA BR Assay Kit (Invitrogen, Massachusetts, US) and stored at -20°C. DNA samples were sequenced on the NovaSeq XP platform with paired-end sequencing, generating high-resolution microbial community profiles. One sample was prefiltered and MgCL2 was added to enrich potential viral sequences [7].

### Sequence Analysis

Adapters and low-quality reads were trimmed using fastp v0.23.4 with default settings [8]. Reads from each sample were assembled using the SPAdes v4.0.0 genome assembler with option “--meta” (metaSPAdes; [9]). Coverage of each contig across all samples was calculated using fairy v0.5.7 [10]. Metagenomic bins were then inferred from bins for each sample, using coverages across all samples, with MetaBAT2 v2.17 with a minimum contig length set to 2kb [11]. Bin quality was assessed using CheckM2 v1.0.1 [12].

Bins were annotated with prokka v1.14.6 [13] and eggnogmapper v 2.1.12 [14]. We predicted the maximum growth rate of each bin using gRodon v 2.4.0 [15]. Taxonomy was assigned to each bin using gtdb-tk v2.1.1 [16]. We used CoverM v0.7.0 to assess bin abundances across samples [17], and bin relative abundances were mclr transformed using the SPRING v1.0.4 R package [18].

We also ran both prokka v1.14.6 [13] (with option metagenome) and gRodon v2.4.0 [19] (with option metagenome_v2) to obtain bulk growth rate predictions for each microbial community. We used sylph v0.8.0 for rapid community-level taxonomic profiling [20] and the R package vegan v2.6-8 for NMDS analysis [21].

## Results

### Community Composition

In general, taxonomic abundances across sample dates and sites remained relatively constant (Fig 1a-d), though samples tended to group by date and by site within dates in their composition (Fig 1e). We noted that early-season samples (July, August) had a higher proportion of *Rhodobacteriales*, whereas later season samples (September) tended to have a higher proportion of *Pelagibacteriales* and *Flavobacteriales* (Fig 1c). One sample, the lone sample taken from site “R” representing water sampled directly from the shore of the Upper New York Bay rather than from our two tidal lagoons, had a distinct taxonomic composition with a higher proportion of *Pelagibacteriales* and a low proportion of both *Rhodobacteriales* and *Flavobacteriales*.

**Figure 1.**
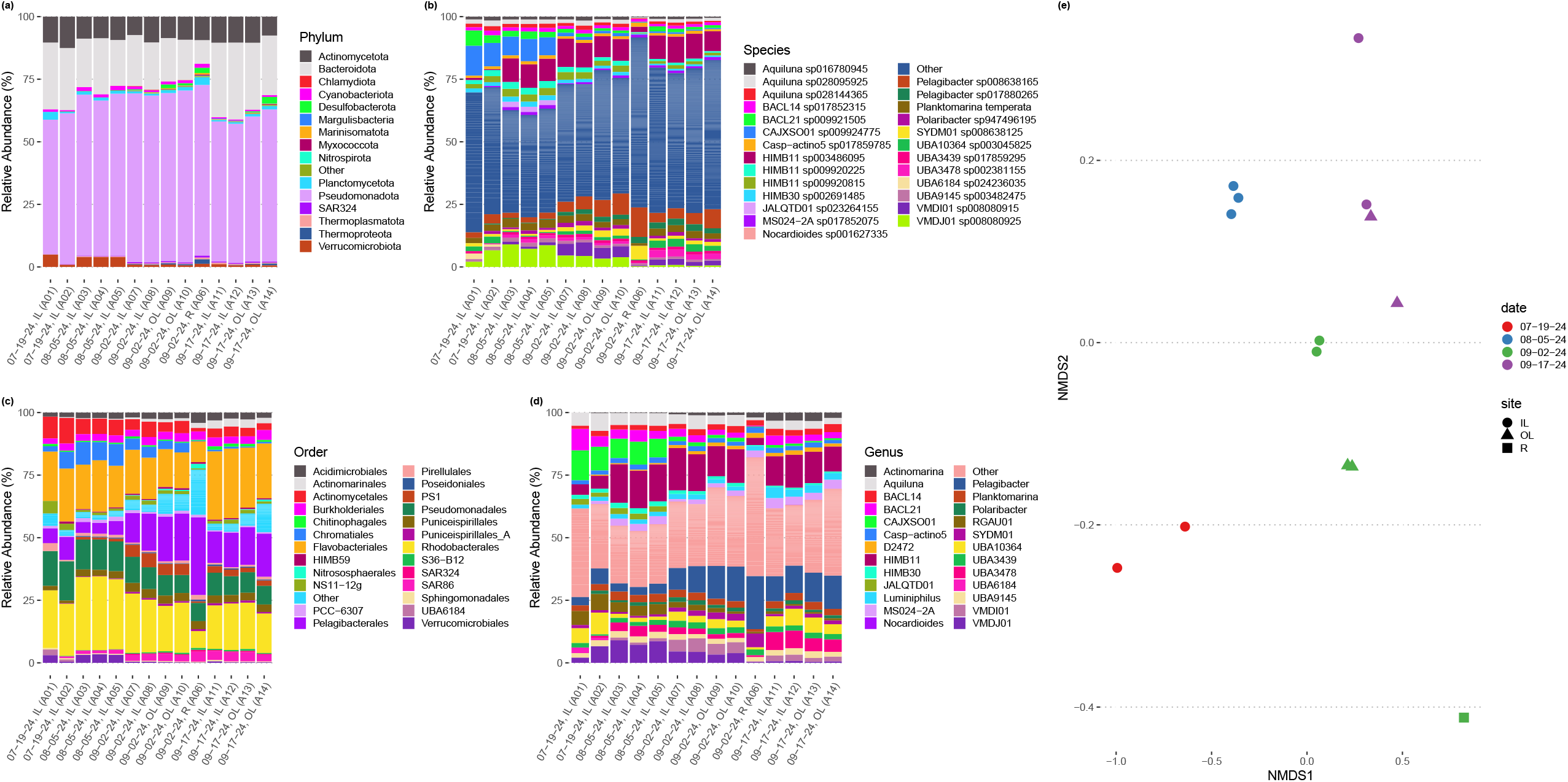
Taxonomic composition of tidal lagoons over the course of a summer. (a-d) Relative abundance of taxonomic groups in each metagenomic sample at various levels of taxonomic resolution. (e) Two dimensional non-metric multidimensional scaling plot of species-level taxonomic composition across our samples groups sampled by site and date. Taxonomic composition inferred by sylph [17].

### Reconstructed Bins

We obtained 1016 total bins, 129 of which were determined to be high quality with less than 5% contamination and being over 90% complete with the total number of contigs ranging from 8-692 and the average contig length ranging from 4,785-372,700bp (S1 Table; [22]). Another 366 were determined to be of medium quality (<10% contamination,>50% completeness). All bins have annotations, including trait data, but we restrict our discussion of results to our high-quality bins. Our high-quality bins span 10 phyla and at least 45 genera. A total of 16 high-quality bins could not be confidently assigned to a known genus and our lone bin from the *Chlamydiota* could not be assigned to a known family, potentially representing novel diversity at these taxonomic levels.

### Trait Data

These bins have diverse functional content on the basis of assigned gene families, with bins from the same phylum typically having a similar number of functional gene assignments but with a great deal of variation both within and between phyla (Fig. 2). Notably, our bins span a range of growth classes, including slow-growth classes that are often missed by isolation-based methods (Fig 3a-b; [15]).

**Figure 2.**
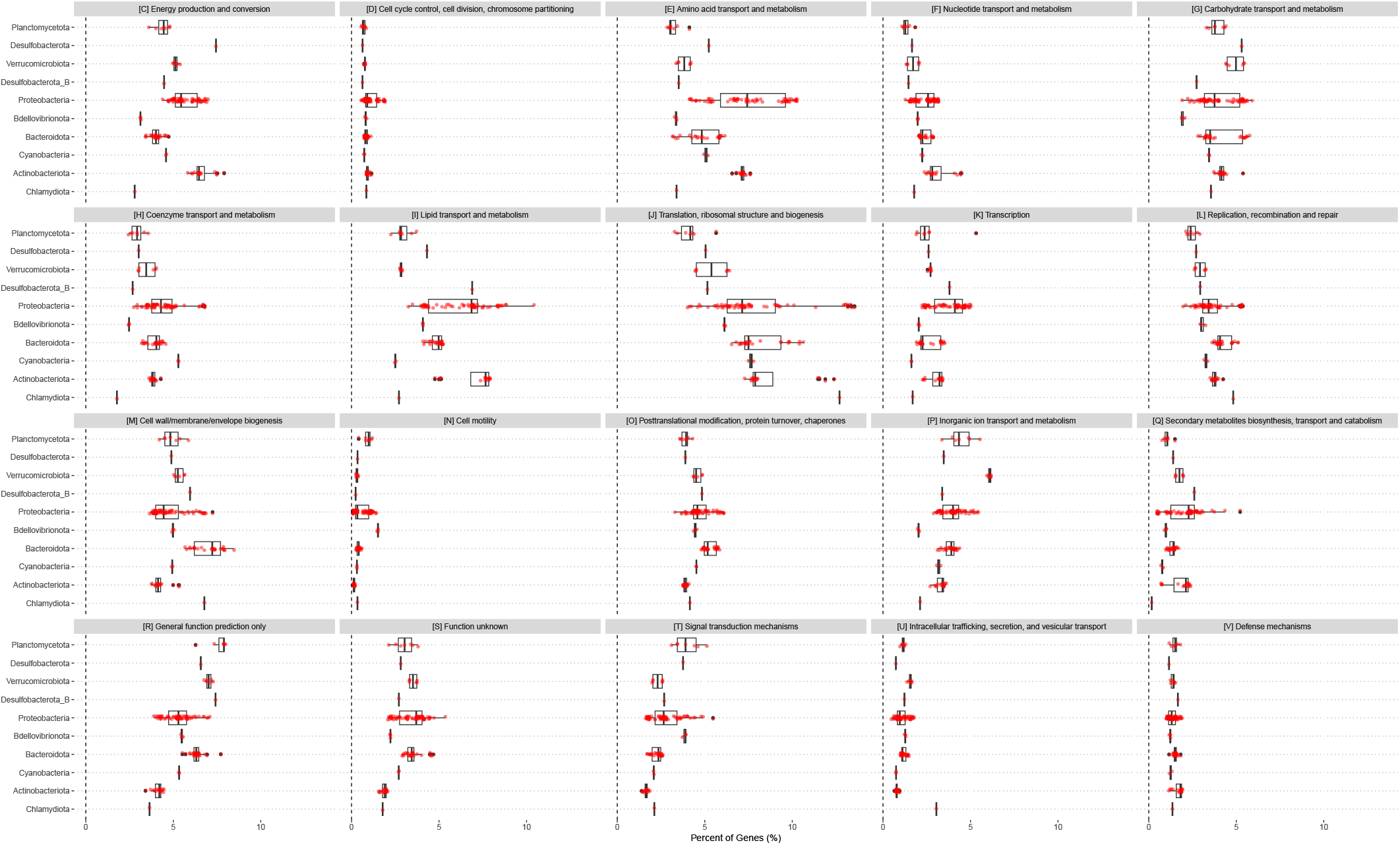
Functional content of high-quality metagenomic bins. Each point is one bin, with the percent of genes belonging to a particular functional class in that bin represented on the x-axis. Functional classifications given by eggnogmapper [11].

**Figure 3.**
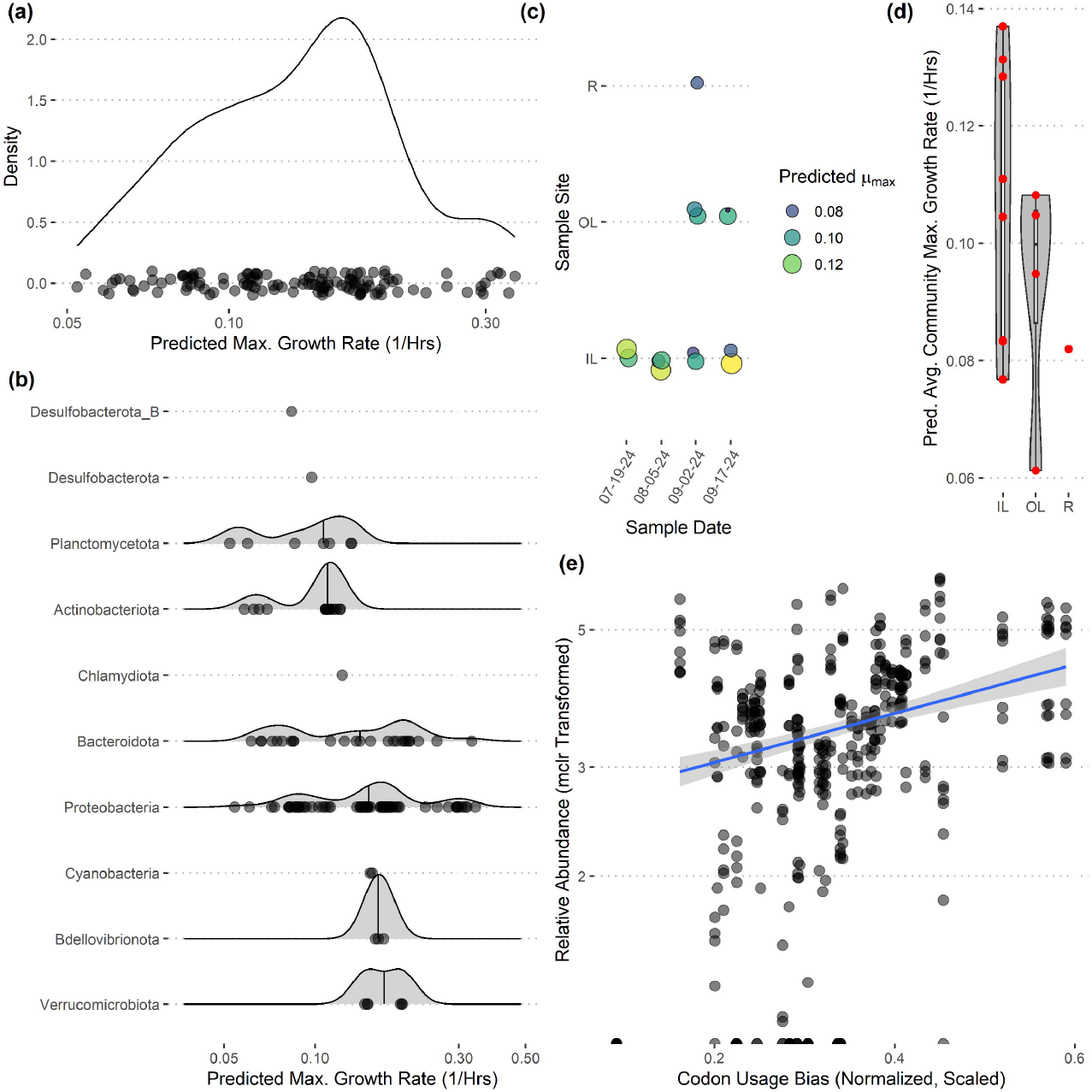
Predicted maximum growth rates for metagenomes and metagenomic bins. (a-b) Distribution of predicted maximum growth rates for metagenomic bins assuming a reference temperature of 25C. (c-d) Predicted community-wide average maximum growth rates for each metagenomic sample. (e) The relative abundances of individual bins across inner-lagoon samples show a positive association with the codon usage bias of each bin. All growth rates and codon usage bias inferred using gRodon [12,16].

Community-wide average maximum growth rate predictions varied across sample sites, with inner lagoon samples seeming to have higher growth rates, though our sample sizes were insufficient to detect any significant effect of sample site on growth (Fig 3c-d; ANOVA, p>0.39, df=2, F=0.999). Looking across inner-lagoon samples, for which we had the most data, the relative abundance of bins was correlated with that bin’s codon usage bias, which is the basis for our genomic maximum growth rate predictions, indicating that increased genomic growth optimization is correlated with higher relative abundances in these samples (Fig 3e; linear regression, p<1e-16, adjusted r^2^=0.166, coefficient=5.97).

## Discussion

We present a comprehensive baseline metagenomic dataset for the urban tidal lagoons located at Bush Terminal Piers Park in Brooklyn, NY, including 15 shotgun metagenomes and 129 high-quality metagenome-assembled genomes (MAGs) with rich functional and taxonomic annotations.

Our community-level data revealed overwhelmingly stable taxonomic composition despite daily flushing by the tides (Fig 1e), with a pattern of gradual taxonomic succession over the course of the season. In a comparison to water sampled directly from the Upper Bay of New York, both tidal lagoons had distinct taxonomic patterns. Stable differentiation between the lagoons and surrounding waters despite flooding with each tide suggests that either (1) the local environment quickly seeds microbes into these waters (e.g. from the surrounding sediments), or (2) by the time of sampling at low-tide the microbial communities in these waters have responded to changes in local conditions in a predictable diel pattern (e.g., shallower, stagnant conditions with abundant invertebrates present including oysters and crabs). We expect the reality to be some combination of the two. In contrast, we did not see any directional pattern of succession over time in our community-level maximum growth rate predictions (Fig 3c-d), although there may be differentiation across sample sites (not significant, ANOVA, F=0.999).

Our reconstructed MAGs had diverse taxonomic affiliations and functional content. Notably, 16 of our MAGs could not be classified at the genus level to anything in the GTDB v220 taxonomy. These MAGs had a range of predicted growth rates that suggested many would not have been readily captured by short-term culturing approaches that often miss slow-growing organisms (maximum growth rates greater than 0.13 in Fig 3a, corresponding to minimum doubling times longer than 5 hours; [15]). We also captured MAGs that ranged widely in their abundances across samples, with fast-growing MAGs predicted to have the highest relative abundances on average (Fig. 3e). These MAGs varied greatly in the proportion of their coding genome associated with particular functions (Fig 2), suggesting that this library covers of range of ecological niches.

## Conclusions

As a dense, coastal city, NYC serves as a valuable model for understanding how climate change-related extreme weather events and sea level rise will impact complex socio-ecological systems. In particular, New York parks serve as potential sites of both social and physical climate resilience, providing relief from recurring heatwaves and flooding events at the same time they allow for community organizing in areas that have suffered a historic lack of investment. Our metagenomic database for this site provides an important baseline for ongoing studies of the microbial communities of New York City’s parks and waterfront areas.

## Data Availability

Metagenomes are available in SRA under BioProject PRJNA1251010. High-quality bins with annotations and code to generate figures are available on Zenodo [23]. Scripts to run metagenomic analysis available at https://github.com/jlw-ecoevo/bushterminalnyc-metagenomics.

